# Transcriptomics of monarch butterflies (*Danaus plexippus*) reveals strong differential gene expression in response to host plant toxicity, but weak response to parasite infection

**DOI:** 10.1101/618546

**Authors:** Wen-Hao Tan, Tarik Acevedo, Erica V. Harris, Tiffanie Y. Alcaide, James R. Walters, Mark D. Hunter, Nicole M. Gerardo, Jacobus C. de Roode

**Affiliations:** Department of Biology, Emory University, Atlanta, Georgia, 30322, USA; Department of Ecosystem Science and Management, Pennsylvania State University, State College, Pennsylvania, USA; Department of Ecology and Evolutionary Biology, University of Kansas, Lawrence, Kansas, USA; Department of Ecology & Evolutionary Biology, University of Michigan, Ann Arbor, Michigan, USA

**Keywords:** RNAseq, secondary metabolites, cardenolides, immunity, *Asclepias*, Lepidoptera

## Abstract

Herbivorous insects have evolved many mechanisms to overcome plant chemical defenses, including detoxification and sequestration. Herbivores may also use toxic plants to reduce parasite infection. Plant toxins could directly interfere with parasites or could enhance endogenous immunity. Alternatively, plant toxins could favor down-regulation of endogenous immunity by providing an alternative (exogenous) defense against parasitism. However, studies on genome-wide transcriptomic responses to plant defenses and the interplay between host plant toxicity and parasite infection remain rare. Monarch butterflies (*Danaus plexippus*) are specialist herbivores that feed on milkweeds (*Asclepias* spp.), which contain toxic cardenolides. Monarchs have adapted to cardenolides through multiple resistance mechanisms and can sequester cardenolides to defend against bird predators. In addition, high-cardenolide milkweeds confer medicinal effects to monarchs against a specialist protozoan parasite (*Ophryocystis elektroscirrha*). We used this system to study the interplay between the effects of plant toxicity and parasite infection on global gene expression. Our results demonstrate that monarch larvae differentially express several hundred genes when feeding on *A. curassavica* and *A. incarnata*, two species that are similar in nutritional content but differ substantially in cardenolide concentrations. These differentially expressed genes include genes within multiple families of canonical insect detoxification genes, suggesting that they play a role in monarch toxin resistance and sequestration. Interestingly, we found little transcriptional response to infection. However, parasite growth was reduced in monarchs reared on *A. curassavica*, and in these monarchs, a small number of immune genes were down-regulated, consistent with the hypothesis that medicinal plants can reduce reliance on endogenous immunity.

## 1 INTRODUCTION

Plants and herbivorous insects have often been used for studying coevolutionary arms races within the framework of chemical ecology (Rosenthal & Berenbaum, 1991). Plants have evolved many forms of defense against herbivores, such as the production of toxic secondary chemicals, and herbivorous insects have evolved mechanisms to overcome such plant defenses (Schoonhoven, van Loon, & Dicke, 2005). These mechanisms include contact avoidance, rapid excretion, sequestration, enzymatic detoxification, and target site mutation (Després, David, & Gallet, 2007). Because host plants species vary in their secondary chemicals, herbivorous insects often utilize different mechanisms when feeding on different plants. For instance, milkweed aphids (*Aphid nerii*) differentially express several canonical insect detoxification genes, including genes encoding Cytochrome P450s (CYP450s), UDP glucuronosyltransferases (UGTs), ATP-binding cassette transporters (ABC transporters), and Glutathione S-transferases (GSTs), when feeding on milkweed species that differ in toxicity (Birnbaum, Rinker, Gerardo, & Abbot, 2017). *Heliconius melpomene* also differentially express UGTs and GSTs when feeding on *Passiflora* species that differ in cyanogen content (Yu, Fang, Zhang, & Jiggins, 2016). Herbivorous insects that feed on widely differing plant families have the additional complication that they may encounter an expanded range of phytochemicals, favoring plastic responses. Indeed, previous work has shown that the Swedish comma butterfly (*Polygonia c-album*) differentially expresses digestion- and detoxification-related genes, as well as genes encoding membrane transporters and cuticular proteins, when feeding on different host plant families (Celorio-Mancera et al., 2013).

While the ability to avoid, resist or excrete toxic chemicals has been selected in many taxa, many insects have also evolved the ability to sequester secondary chemicals into their own tissues, thereby protecting themselves against their own natural enemies (Opitz & Müller, 2009). For example, in Lepidoptera (reviewed in Nishida, 2002), some swallowtail butterflies sequester aristolochic acid from their host plants to deter vertebrate predators (Uésugi, 2010); buckeye butterflies (*Junonia coenia*) sequester iridoid glycosides (IGs), which deter invertebrate predators (Dyer & Bowers, 1996; Theodoratus & Bowers, 1999); and tiger moths (*Grammia incorrupta*) sequester pyrrolizidine alkaloids, which defend them against parasitoids (Singer, Mace, & Bernays, 2009). In addition to the direct effects of sequestered chemicals on anti-predator and –parasite defense, phytochemicals can also indirectly affect parasites by modulating the host immune system (Lampert, 2012). Depending on the particular chemicals and parasites, toxin sequestration may reduce, enhance, or have no effect on anti-parasite immunity. For instance, all three scenarios have been shown in herbivores that sequester IGs. *Junonia coenia* exhibits reduced immunity (measured by the melanization response) when feeding on *Plantago lanceolata*, a plant species with greater concentrations of IGs, than when feeding on *P. major*, a less toxic host plant (Smilanich, Dyer, Chambers, & Bowers, 2009). In contrast, in this same system, feeding on the more toxic plant enhances anti-viral defenses (Smilanich et al., 2017). *Melitaea cinxia* shows enhanced immunity when feeding on *Plantago lanceolata* strains with higher IG concentration (Laurentz et al., 2012), but in *Grammia incorrupta*, a moth species that also feeds on IG-containing plants, IG concentration does not appear to affect immune responses (Smilanich, Vargas, Dyer, & Bowers, 2011).

As described above, phytochemicals pose both challenges and benefits for herbivorous insects, and the ecological interactions and evolutionary relationships between plants and herbivorous insects have been studied extensively. However, studies of genome-wide transcriptomic responses to plant defenses, which provide insight into the simultaneous effects of toxins on detoxification, sequestration, and immune systems, remain rare (Celorio-Mancera et al., 2013; Vogel, Musser, & Celorio-Mancera, 2014). Even for herbivorous insect species with genomic and transcriptomic information available, transcriptomic research has rarely focused on herbivore-plant interactions (Vogel et al., 2014).

Here, we provide a transcriptomics-based analysis of parasite-infected and –uninfected monarch butterflies (*Danaus plexippus*) feeding on different host plant species. Monarch butterflies are a prominent example of sequestration and aposematism (Agrawal, Petschenka, Bingham, Weber, & Rasmann, 2012). Monarchs are specialist herbivores on milkweeds (mostly *Asclepias* spp.), but these plants vary widely in their toxicity, measured predominantly as the concentration and composition of cardenolides (Agrawal et al., 2012). Cardenolides are steroids that are toxic to most animals because they inhibit the essential enzyme Na^+^/K^+^-ATPase that is responsible for maintaining membrane potentials (Agrawal et al., 2012). Monarchs and other herbivorous insects specializing on cardenolide-containing plants have convergently evolved amino acid substitutions on the target site of the toxins that decrease binding affinity (Dobler, Dalla, Wagschal, & Agrawal, 2012; Zhen, Aardema, Medina, Schumer, & Andolfatto, 2012). Target site insensitivity largely enhances monarch resistance to cardenolides, but they are not completely resistant to cardenolides (Agrawal et al., 2012; Petschenka, Offe, & Dobler, 2012). There are fitness costs, including reduced larval survival and adult lifespan, for monarchs feeding on milkweed species with high cardenolide concentration or toxicity (Agrawal, 2005; Malcolm, 1994; Tao, Hoang, Hunter, & de Roode, 2016; Zalucki, Brower, & Alonso-M, 2001; Zalucki, Brower, & Malcolm, 1990; Zalucki & Brower, 1992). Despite these costs, monarchs have evolved the ability to sequester cardenolides into their own tissues, which, coupled with bright warning coloration, deters bird predators (Brower, Ryerson, Coppinger, & Susan, 1968). In addition to the anti-predator protection provided by milkweeds, high-cardenolide milkweeds also provide protection against the common specialist parasite *Ophryocystis elektroscirrha* (de Roode, Pedersen, Hunter, & Altizer, 2008; Sternberg et al., 2012). Monarchs become infected with this parasite during their larval stage when ingesting parasite spores (Mclaughlin & Myers, 1970), but feeding on milkweeds with greater concentrations of cardenolides results in lower parasite infection, growth and virulence (de Roode, Pedersen, et al., 2008; de Roode, Rarick, Mongue, Gerardo, & Hunter, 2011; Gowler, Leon, Hunter, & de Roode, 2015; Lefèvre, Oliver, Hunter, & de Roode, 2010; Sternberg, de Roode, & Hunter, 2015; Sternberg et al., 2012; Tan, Tao, Hoang, Hunter, & de Roode, 2018; Tao, Gowler, Ahmad, Hunter, & de Roode, 2015; Tao, Hoang, et al., 2016). At present, however, it remains unclear how cardenolides, parasites, and the monarch’s immune system interact. On the one hand, it is possible that cardenolides directly interfere with parasites. This could result in a down-regulation of immune responses, as these chemicals would fulfill the same role as anti-parasitic immunity. Alternatively, cardenolides could stimulate the monarch immune system and thus enhance immune responses against parasites. Therefore, monarchs provide an excellent model to study how detoxification, toxin sequestration, and immunity interact in a system with a known association between phytochemicals and disease resistance.

In this study, we assess differential gene expression between monarch larvae feeding on the low-cardenolide *A. incarnata* and the high-cardenolide *A. curassavica* when infected or uninfected with the specialist parasite *O. elektroscirrha*. Specifically, we performed RNA-Seq on two tissue types of parasite-infected and uninfected larvae fed with either plant species. In addition, we quantified parasite resistance of the same batch of larvae and measured foliar cardenolide concentration in the same batch of milkweeds. While we found a limited transcriptional response to parasite infection, our results reveal a large number of genes that are differentially expressed in monarchs reared on the two milkweed species, including the down-regulation of four immune genes when fed on the high-cardenolide *A. curassavica*.

## 2 MATERIALS AND METHODS

### 2.1 Monarchs, milkweeds, and parasites

Monarch butterflies in this study were obtained from a lab-reared, outcrossed lineage generated from wild-caught migratory monarchs collected in St. Marks, Florida, USA. The parasite clone (C_1_-E_25_-P_3_) was isolated from an infected, wild-caught monarch from the same population. We used two species of milkweed in this study: *A. incarnata* and *A. curassavica*.

These two species were chosen because they are similar in nutrient content but differ substantially in their level of cardenolides (toxic, secondary compounds)(Tao, Ahmad, de Roode, & Hunter, 2016); concentrations in *A. curassavica* are generally at least 10-fold higher than are those in *A. incarnata*. As a consequence, the milkweeds have been shown repeatedly to differentially affect monarch resistance to parasitism, with *A. curassavica* reducing parasite infection, growth, and virulence relative to *A. incarnata* (de Roode, Pedersen, et al., 2008; de Roode et al., 2011; Lefèvre et al., 2010; Sternberg et al., 2015, 2012; Tao et al., 2015; Tao, Hoang, et al., 2016). Milkweed seeds were obtained from Prairie Moon Nursery (Winona, MN, USA). All milkweeds in this study were grown in a greenhouse under natural light conditions with weekly fertilization (Jack’s 20-10-20 from JR Peters Inc. Allentown, PA, USA).

### 2.2 Experimental design and sample collection

We used second instar larvae for transcriptome sequencing because larvae most likely become infected with *O. elektroscirrha* during early instars under natural conditions, through either vertical or horizontal transmissions (Altizer, Oberhauser, & Geurts, 2004; de Roode, Chi, Rarick, & Altizer, 2009). We could not use first instars due to size limitations. Also, second instar larvae sequester the highest amounts of cardenolides relative to their body mass (Jones, Peschenka, Flacht, & Agrawal, 2019). Upon hatching, we reared larvae individually in Petri dishes on cuttings from different plants of either *A. incarnata* or *A. curassavica*. We inoculated second instar larvae by adding ten parasite spores to an 8-mm diameter leaf disk taken from the milkweed species upon which they had been feeding, following an established protocol (de Roode, Yates, Altizer, & Roode, 2008). Uninfected controls received leaf disks without spores. After larvae consumed their entire leaf disk, they were provided leaves of the same milkweed species *ad libitum.* Eighteen to twenty-four hours after parasite inoculation, we placed larvae in RNAlater and stored them at 4°C. We dissected all larvae within four days of collection. We separated the entire digestive tract (hereafter, gut) and the remaining body (hereafter, body) and put the samples into separate tubes with RNAlater. We stored these samples at −80 °C. Sample sizes for each treatment group and tissue type are provided in supplemental information Table S1.

We reared another subset of parasite-infected and uninfected larvae to adulthood on each plant species to quantify parasite resistance (N = 9-17 per treatment group). After parasite inoculation, larvae were transferred to individual rearing cups (473 mL) and fed leaves from either *A. curassavica* or *A. incarnata*. After pupation, pupae were placed in a laboratory room maintained at 25 °C under 14/10h L/D cycle. After eclosion, adults were placed in 8.9 × 8.9 cm glassine envelopes without a food source at 12 °C under 14/10h L/D cycle. Parasite load was quantified using a vortexing protocol described in de Roode et al., 2008. Normality and variance homogeneity were checked with the Shapiro-Wilk normality test and Fligner-Killeen test. Parasite spore load data were analyzed using a two-sample t-test. All analyses were performed in R version 3.5.2 (R Core Team, 2018).

### 2.3 Chemical analyses

We collected two types of samples for chemical analyses: milkweed foliage and larval frass. We collected foliage samples to confirm the differences in total cardenolide concentration between the two species. In addition, we collected larval frass to compare the differences between cardenolide composition before and after larval digestion. Foliage samples of the two plant species (N = 11-12 individual plants per species) were collected on the same day that we performed parasite inoculations. One leaf from the fourth leaf pair on each plant was chosen. Six leaf disks (424 mm^2^ total) were taken with a paper hole punch from one side of the leaf and placed immediately into a 1 mL collection tube with cold methanol. Another six identical leaf disks were taken from the opposite side of the same leaf to measure sample dry mass. Frass samples, each from an individual larva, were collected from another subset of second instar larvae that were reared from hatchlings on *A. curassavica* (N = 17). For this analysis, we focused on *A. curassavica* only because *A. incarnata* foliage contains very few cardenolides. Frass samples for each individual were collected for 24 hours during the second instar. Frass samples were collected into 1 mL collection tubes with cold methanol on the day of frash production. Total cardenolide concentrations and cardenolide compositions were analyzed using reverse-phase ultra-performance liquid chromatography (UPLC; Waters Inc., Milford, MA, USA) following established methods (Tao et al., 2015). The absorbance spectra were recorded from 200 to 300 nm with digitoxin used as an internal standard. Under reverse-phase UPLC, cardenolide retention time decreases as polarity increases. For the plant samples, we analyzed the difference in total cardenolide concentration between the two species. Normality and variance homogeneity were checked with the Shapiro-Wilk normality test and Fligner-Killeen test. Cardenolide data were analyzed using a Mann–Whitney U test due to violation of assumptions of normality and variance homogeneity. All analyses were performed in R version 3.5.2 (R Core Team, 2018). We assessed the differences in cardenolide compositions by comparing the cardenolide peaks between the two sample types.

### 2.4 RNA extraction, library preparation, and sequencing

We extracted total RNA from either gut or body tissues using the RNeasy RNA mini extraction kit (Qiagen) following the manufacturer’s protocol. The quality and quantity of RNA samples were assessed using a nanodrop and bioanalyzer. Total RNA was sent to BGI (Beijing Genomics Institute, Hong Kong) for library preparation and sequencing. We sequenced the two tissue types (gut and body separately) of infected and uninfected larvae fed with either *A. incarnata* or *A. curassavica*, with 3-4 biological replicates per treatment (see supplemental information Table S1). We performed 50 bp single-end sequencing with a sequencing depth of 20M reads per sample using the BGIseq-500 platform.

### 2.5 Transcriptome assembly

We checked the quality of RNA-seq reads using FastQC (Andrews, 2010) and compiled across samples using MultiQC (Ewels, Magnusson, Lundin, & Käller, 2016). Sequence quality was consistently high across positions (see supplemental information Fig. S1), so we proceeded without trimming. RNA-seq reads for each sample were mapped to the monarch reference genome (Zhan, Merlin, Boore, & Reppert, 2011) using STAR ver 2.5.2b (Dobin et al., 2013) and checked for alignment statistics. There were two samples that had low quality; one of them had a very low quantity of reads and the other had a very low mapping rate. Given that these two samples were from different individuals, we removed four samples (i.e., both tissue types of the same individual) from our analyses. We obtained the number of reads mapped to each gene from STAR and compiled them across samples as a count matrix.

### 2.6 Differential gene expression analysis

Differential gene expression analysis was performed using the R Bioconductor package edgeR version 3.24.3 (Robinson, McCarthy, & Smyth, 2009). We performed separate analyses on the two tissue types. We removed genes without any counts across samples from our analyses. We normalized the library sizes across samples using the trimmed mean of M-values (TMM) normalization. We performed differential gene expression analyses using negative binomial generalized linear models (GLMs). We created design matrices for GLM with infection treatment and plant species as factors, estimated dispersion parameters, and fitted the models. We addressed specific questions of interest by setting coefficient contrasts to compare between different treatment groups. First, we compared gene expression between all infected and all uninfected larvae to examine the overall impacts of parasite infection. We then compared gene expression between infected and uninfected larvae reared on the two milkweeds species separately to examine plant-specific effects. Next, we compared gene expression between larvae fed with *A. incarnata* and *A. curassavica*; given that we found almost no differences between infected and uninfected groups, we combined them for this comparison. The Benjamini-Hochberg method (Benjamini & Hochberg, 1995) was used to account for multiple hypothesis testing and to calculate adjusted p-values. We visualized the results through heatmaps with hierarchical clustering, MA plots, and volcano plots generated using the R package edgeR version 3.24.3 (Robinson et al., 2009) and gplots version 3.0.1 (Warnes et al., 2016). All analyses were performed in R version 3.5.2 (R Core Team, 2018).

### 2.7 Examine specific gene sets of interest

Given that we were specifically interested in genes that function in immunity and detoxification, we examined if canonical immune genes and detoxification genes were differentially expressed among treatment groups. We obtained a full set of annotated monarch immune genes published by the *Heliconius* Genome Consortium (2012), which included a set of annotated (*Heliconius*) immune genes and their orthologs in several species, including monarchs. The monarch orthologs listed in this published dataset were based on a previous version of monarch genome annotation (OGS1.0), so we updated this full set of immune genes to the latest version of gene annotation (OGS2.0) using information provided in Monarch Base (Zhan & Reppert, 2013). This updated monarch immune gene set contains 114 genes belonging to the functional classes of recognition, signaling, modulation, and effector (see supplemental information Table S2). For detoxification genes, similar to a previous study on another milkweed-feeding insect (Birnbaum et al., 2017), we focused on four canonical gene families: Cytochrome P450s (CYP450s), UDP glucuronosyltransferases (UGTs), ATP-binding cassette transporters (ABC transporters), and Glutathione S-transferases (GSTs). We obtained those annotated detoxification genes from Monarch Base (Zhan & Reppert, 2013). We examined each set of our significantly differentially expressed genes to obtain the number of immune and detoxification genes within them. For all the significantly differentially expressed detoxification genes, we performed BLAST searches against two other Lepidopteran species (*Bombyx mori* and *Heliconius melpomene*) via the EnsemblMetazoa database (https://metazoa.ensembl.org/) to verify that their top hit paralogs also have the same putative detoxification function.

### 2.8 Gene ontology enrichment analysis

Functional annotations and Gene Ontology (GO) term assignments for all protein-coding genes in the genome were generated using PANNZER2 (Törönen, Medlar, & Holm, 2018), with protein sequences obtained from Monarch Base, using default parameters. We created a custom annotation package for our organism using AnnotationForge (Carlson & Pages, 2018). We performed GO-term enrichment analyses on differentially expressed genes using ClusterProfiler (Yu, Wang, Han, & He, 2012) with default p-value and q-value cutoff thresholds. The “gene universe” included all genes that were expressed in our RNA-Seq dataset. The Benjamini-Hochberg method (Benjamini & Hochberg, 1995) was used to account for multiple hypothesis testing and to calculate the adjusted p-values. We included all three ontology groups in our analyses: biological process (BP), molecular function (MF), and cellular components (CC). We visualized the enrichment results by dotplots using ClusterProfiler (Yu et al., 2012)

## 3 RESULTS

### 3.1 Plant chemistry and parasite resistance

We confirmed previous findings that the two milkweed species differ greatly in cardenolide concentration and differentially affect monarch resistance to parasitism. Total cardenolide concentration of *A. curassavica* foliage was 95-fold higher than that of *A. incarnata* foliage (Fig. 1A; *W* = 0, *P* < 0.0001), and butterflies reared on *A. curassavica* experienced significantly lower parasite spore load than those fed with *A. incarnata* (Fig. 1B; *t* = 3.39, *df* = 19, *P* = 0.003). None of the uninoculated monarchs became infected (N = 9 for *A. incarnata* and N = 17 for *A. curassavica*). When comparing the cardenolide composition of *A. curassavica* foliage and the frass from larvae feeding on *A. curassavica*, we found that they differed greatly in composition (Fig. 2). Specifically, out of a total of 22 unique cardenolides (i.e., individual bars in Fig. 2), only four occurred in both foliage and frass; eight cardenolides were exclusively found in foliage, and nine were exclusively found in frass. Additionally, there were more polar cardenolides in frass than in foliage, as indicated by lower retention times relative to a digitoxin internal standard (Fig. 2).

**Figure 1.**
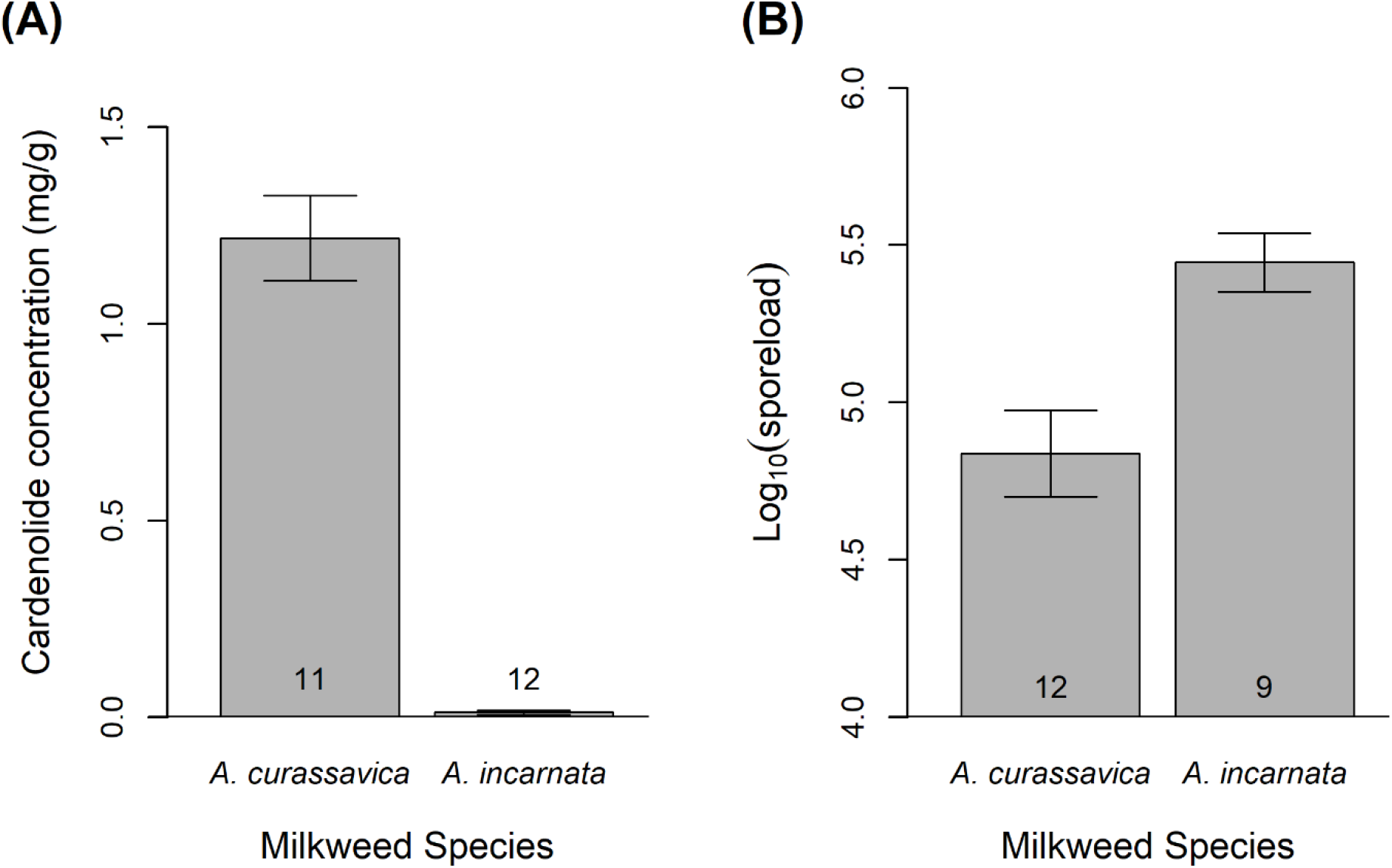
Differences in foliar cardenolide concentration and monarch parasite resistance between the two milkweed species, *A. curassavica* and *A. incarnata*. (A) Total cardenolide concentraion of foliage. (B) The effect of milkweed species on parasite spore load in infected monarchs. Data represent mean ±1 SEM. Sample sizes are reported on each bar.

**Figure 2.**
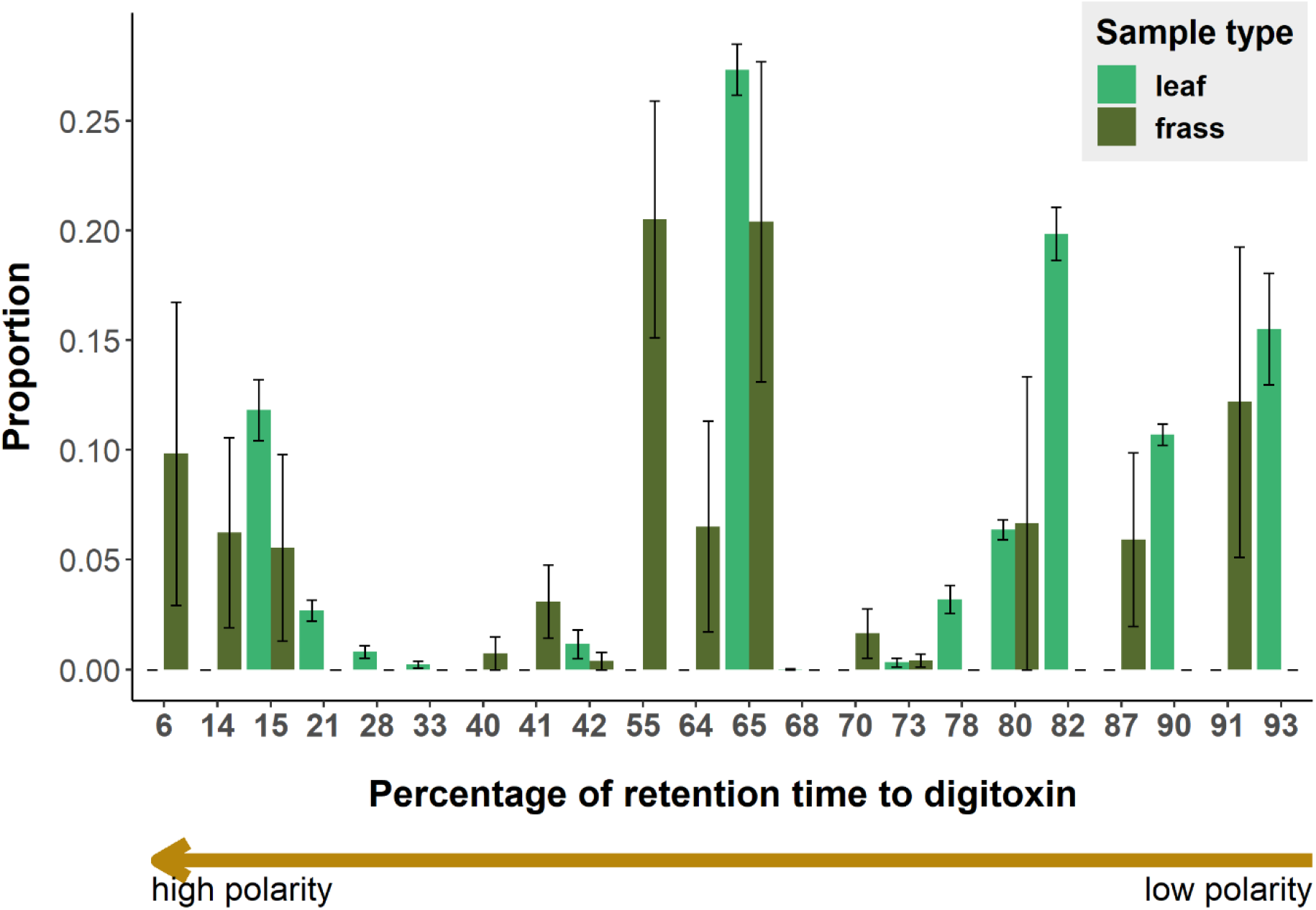
Cardenolide composition of *A. curassavica* foliage and frass produced by larvae fed with *A. curassavica*. The X-axis represents the percentage of retention time relative to a digitoxin internal standard in UPLC. Bars represent individual cardenolides. The Y-axis represents the proportion of the individual cardenolide within each sample. Data represent the mean ±1 SEM. Sample sizes: N = 11 for foliage samples (each sample was collected from a different individual plant) and N = 17 for frass samples (each sample was collected from a different individual larva). We only focused on *A. curassavica* because *A. incarnata* foliage contains very few cardenolides.

### 3.2 Differential gene expression analysis in relation to parasite infection

We first compared gene expression between all infected and all uninfected larvae to examine the overall effects of parasite infection on gene expression. Surprisingly, in both gut and body tissues, we found that no genes were significantly differentially expressed (Fig. 3-4, Table 1). Next, we compared gene expression between infected and uninfected larvae reared on the two milkweed species separately to examine plant-specific effects. Again, we found almost no response to parasite infection (Table 1). For the larvae fed with *A. incarnata*, only one gene was significantly up-regulated in the gut in the infected group when compared to the uninfected group: a cytochrome P450 gene (DPOGS205609). For the larvae fed with *A. curassavica*, only two genes were significantly down-regulated in the body in the infected group: an acid digestive lipase (DPOGS211626) and a carboxypeptidase (DPOGS211663). Overall, we found extremely few differentially expressed genes between infected and uninfected larvae regardless of tissue type or host plant, and none of those that were significantly differentially expressed were canonical immune genes.

**Figure 3.**
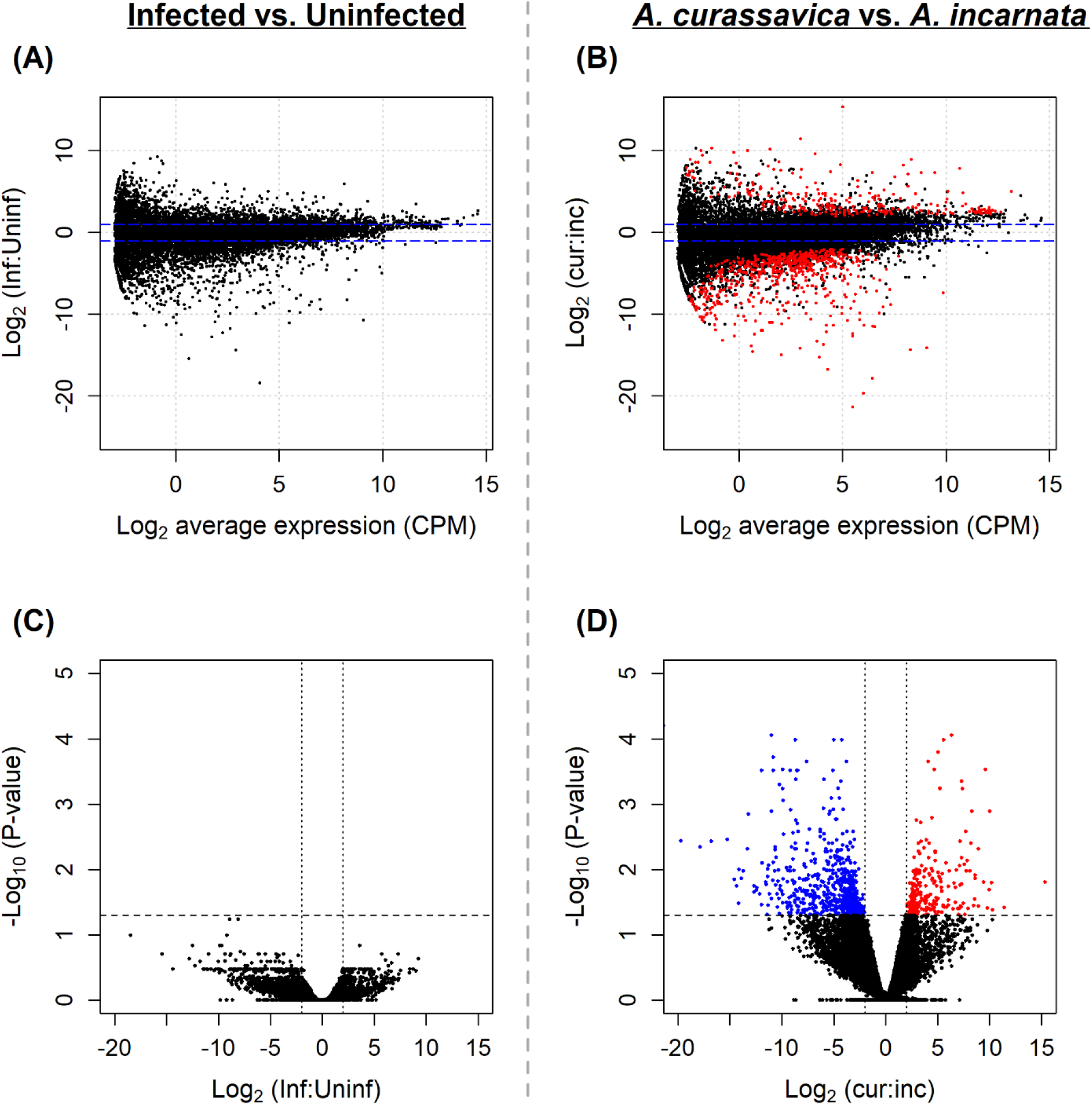
Patterns of differential gene expression in gut tissue. (A) and (C): expression differences between infected and uninfected larvae. A positive fold change indicates up-regulation in infected larvae. (B) and (D): expression differences between larvae fed with *A. curassavica* and *A. incarnata*. A positive fold change indicates up-regulation in larvae fed with *A. curassavica*. (A) and (B): MA plots. Dotted horizontal lines indicate ± 1-fold change. (C) and (D): volcano plots. Dotted horizontal lines indicate p-value thresholds. Dotted vertical lines indicate ± 2-fold change. Blue dots represent significantly down-regulated genes; red dots represent significantly up-regulated genes.

**Figure 4.**
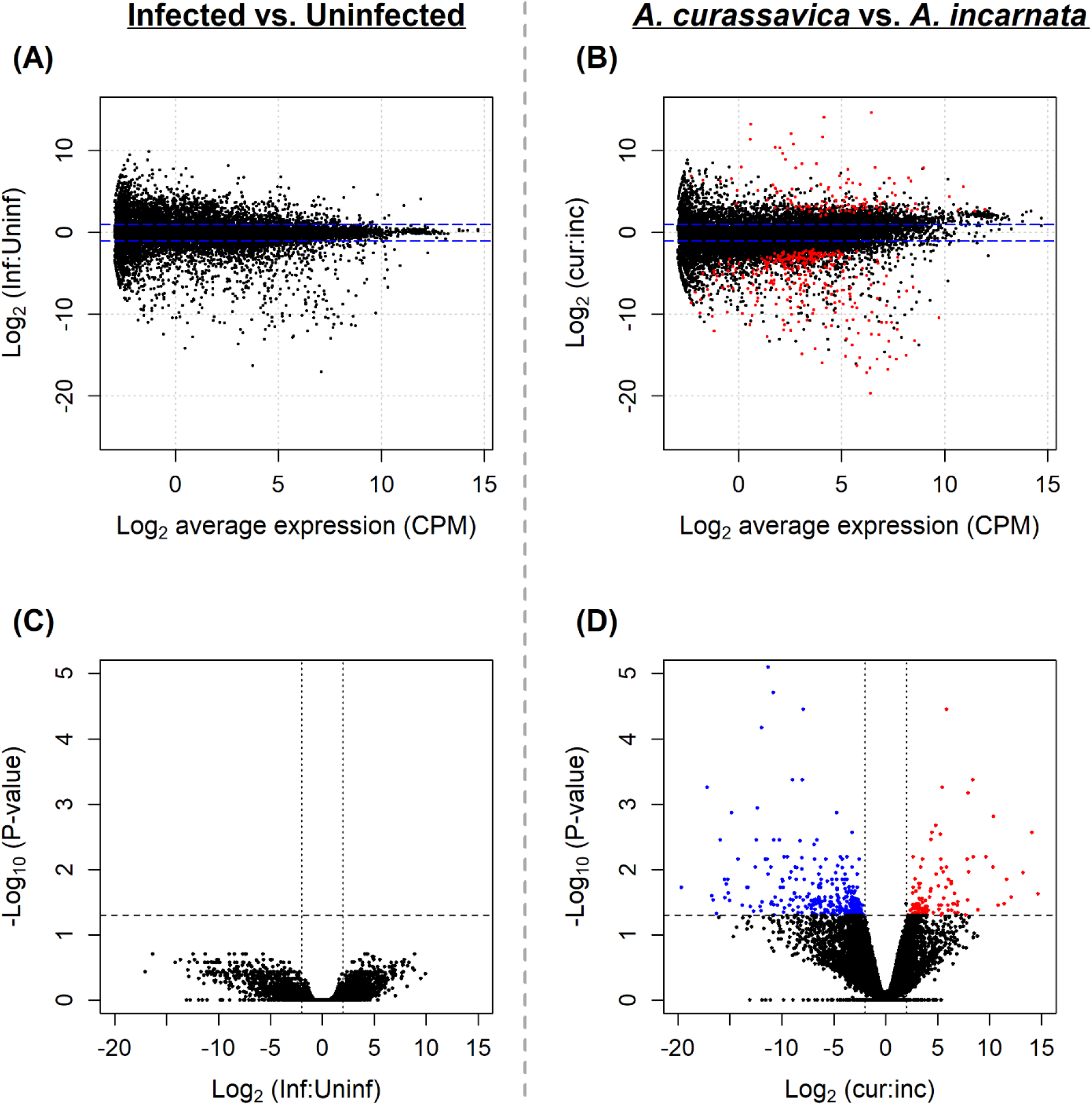
Patterns of differential gene expression in body tissue. (A) and (C): expression differences between infected and uninfected larvae. A positive fold change indicates up-regulation in infected larvae. (B) and (D): expression differences between larvae fed with *A. curassavica* and *A. incarnata*. A positive fold change indicates up-regulation in larvae fed with *A. curassavica*. (A) and (B): MA plots. Dotted horizontal lines indicate ± 1-fold change. (C) and (D): volcano plots. Dotted horizontal lines indicate p-value thresholds. Dotted vertical lines indicate ± 2-fold change. Blue dots represent significantly down-regulated genes; red dots represent significantly up-regulated genes.

**Table 1.** Summary of differentially expressed genes. The first two columns denote specific comparisons and the subset of samples used. The last three columns indicate the number of significantly up-regulated and down-regulated genes upon infection, or between those fed with different milkweed species, in either gut tissue or body. First, we compared infected and uninfected larvae in all samples to assess overall transcriptional patterns of parasite infection (*i.e.*, the first row). We then compared infected and uninfected larvae reared on the two milkweed species separately to examine plant-specific effects (*i.e*., the second and third rows). Next, we compared larvae fed with *A. incarnata* and *A. curassavica*. Given that we found almost no differences between infected and uninfected groups, we combined them for this comparison (*i.e.*, the fourth row).

| Factor | Subset | Direction | Gut | Body |
| --- | --- | --- | --- | --- |
| Infection | All | up-regulated in infected | 0 | 0 |
|  |  | down-regulated in infected | 0 | 0 |
| Infection | *A. incarnata* | up-regulated in infected | 1 | 0 |
|  |  | down-regulated in infected | 0 | 0 |
| Infection | *A. curassavica* | up-regulated in infected | 0 | 2 |
|  |  | down-regulated in infected | 0 | 0 |
| Plant | All | increased in *A. curassavica* | 271 | 122 |
|  |  | Increased in *A. incarnata* | 637 | 306 |

### 3.3 Differential gene expression analysis in relation to milkweed diet

We compared gene expression between larvae reared on *A. curassavica* and *A. incarnata.* Given that we found almost no differences in expression between infected and uninfected larvae, we combined them in this comparison between plant species. We found that 908 genes were differentially expressed in the gut and 428 genes were differentially expressed in the body (Fig. 3-4, Table 1). Given that the gut is the place where initial digestion of plant matter happens, we expected the transcriptional patterns to be more distinct between plant diets in gut than in body samples. Indeed, heatmap and hierarchical clustering suggest that individuals are more clustered by plant diet in gut samples than in body samples (Fig. 5). The top 15 up-regulated and top 15 down-regulated genes for the gut and body are listed in Table 2 and Table 3, respectively. In gut tissues, notably, one of the top 15 up-regulated genes when fed with *A. curassavica* is a glutathione S-transferase (DPOGS210488), and another one is a carboxyl esterase (DPOGS204275), both of which are canonical insect detoxification genes and possibly might function in processing cardenolides. Other genes belong to a variety of biological functions, such as digestive processes and membrane-related proteins. Differential expression of digestive and membrane-related genes has also been demonstrated in other insects when feeding on different plant species (Celorio-Mancera et al., 2013).

**Figure 5.**
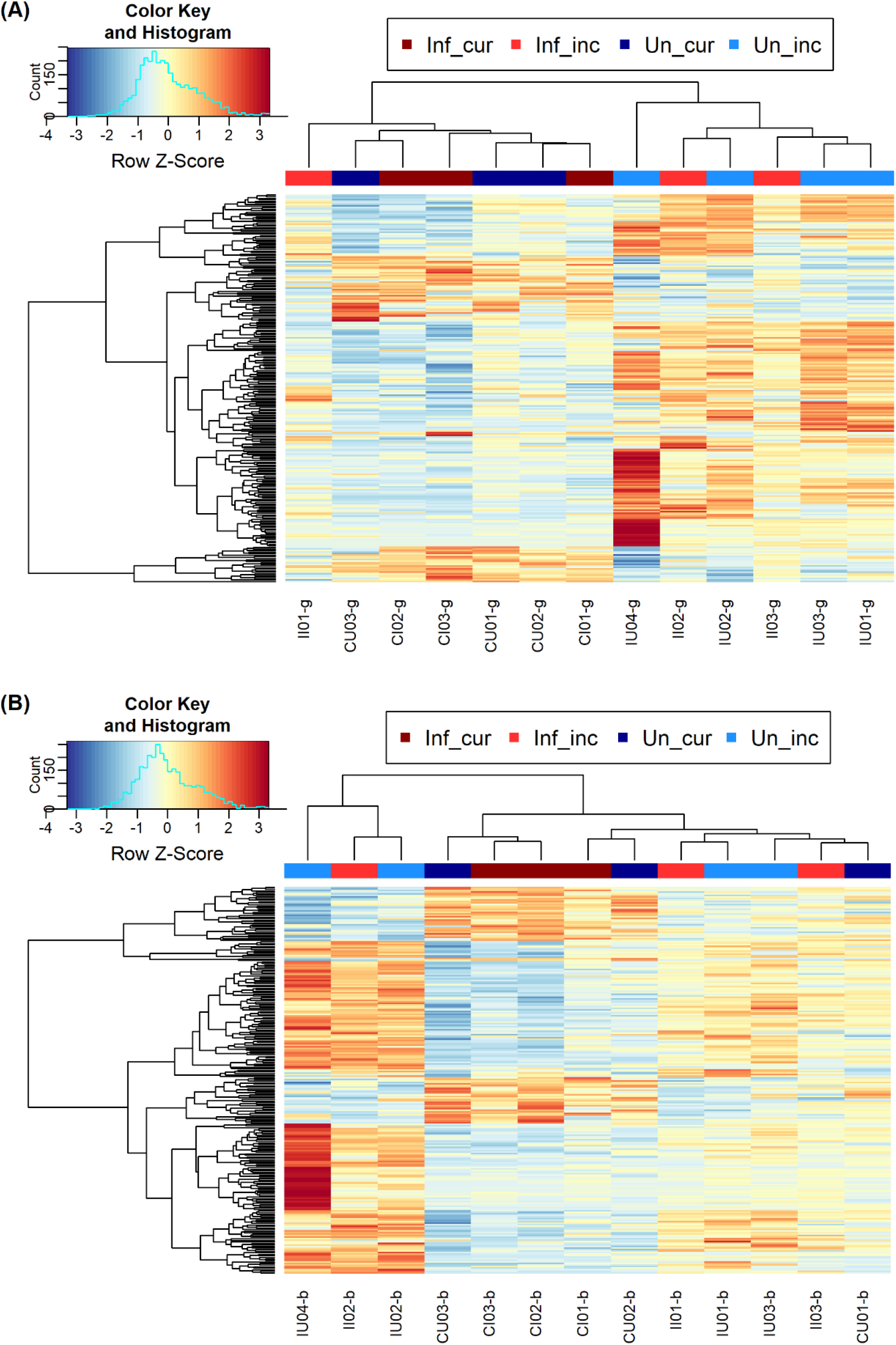
Heatmap and hierarchical clustering of the top 250 differentially expressed genes between larvae fed with *A. curassavica* and *A. incarnata*. (A) The result of gut samples. Hierarchical clustering shows that samples are clustered mostly based on the plant species larvae were fed with. (B) The result of body samples. The clustering patterns are less clear. “Inf_cur” represents infected larvae fed with *A. curassavica*; “Inf_inc” represents infected larvae fed with *A. incarnata*; “Un_cur” represents uninfected larvae fed with *A. curassavica*; “Un_inc” represents uninfected larvae fed with *A. incarnata*.

**Table 2.** List of top 15 differentially expressed genes in gut tissue between larvae fed with *A. curassavica* and *A. incarnata*. The list includes the top 15 genes significantly up-regulated when <u>fed</u> with *A. curassavica* and the top 15 genes significantly up-regulated when fed with *A. incarnata*.

| Gene ID | log<sub>2</sub>FC | logCPM | FDR | Protein |
| --- | --- | --- | --- | --- |
| ***Top 15 up-regulated genes in A. curassavica*** |  |  |  |  |
| DPOGS201344 | 6.372 | 5.747 | 8.896E-05 | Uncharacterized |
| DPOGS202254 | 5.589 | 5.739 | 1.040E-04 | Threonine dehydratase catabolic-like isoform 2 |
| DPOGS215709 | 5.049 | 13.155 | 1.596E-04 | Uncharacterized |
| DPOGS212746 | 4.112 | 10.044 | 2.210E-04 | Uncharacterized |
| DPOGS213427 | 4.699 | 4.654 | 2.947E-04 | Phosphatidyltransferase |
| DPOGS204785 | 9.623 | 3.669 | 2.947E-04 | Carboxypeptidase 4 |
| DPOGS209145 | 7.309 | 6.446 | 4.455E-04 | Uncharacterized |
| DPOGS204275 | 5.239 | 3.825 | 5.752E-04 | Carboxyl/choline esterase |
| DPOGS213104 | 7.410 | 4.420 | 5.799E-04 | Zinc finger protein |
| DPOGS204877 | 5.220 | 7.017 | 5.799E-04 | Uncharacterized |
| DPOGS210488 | 10.030 | -1.820 | 1.296E-03 | Glutathione S-transferase epsilon 4 |
| DPOGS205617 | 8.315 | 4.894 | 1.296E-03 | Gucocerebrosidase |
| DPOGS200701 | 4.470 | 3.245 | 1.614E-03 | Spliceosomal protein |
| DPOGS214834 | 2.985 | 6.014 | 1.746E-03 | Juvenile hormone epoxide hydrolase |
| DPOGS206961 | 3.390 | 6.869 | 1.906E-03 | Fructose 1,6-bisphosphate aldolase |
| ***Top 15 up-regulated genes in A. incarnata*** |  |  |  |  |
| DPOGS213127 | -14.990 | 2.053 | 2.820E-06 | Nuclear receptor GRF |
| DPOGS209249 | -21.366 | 5.499 | 6.322E-05 | Uncharacterized |
| DPOGS205455 | -11.005 | 1.492 | 8.896E-05 | Uncharacterized |
| DPOGS215049 | -8.676 | 3.715 | 1.040E-04 | Peroxidasin-like protein |
| DPOGS214337 | -4.961 | 2.053 | 1.040E-04 | Dystrophin |
| DPOGS206024 | -4.189 | 4.407 | 1.040E-04 | Uncharacterized |
| DPOGS205589 | -10.789 | 5.246 | 1.909E-04 | Hormone receptor 3C |
| DPOGS215508 | -3.738 | 3.000 | 2.210E-04 | Uncharacterized |
| DPOGS210943 | -7.584 | 5.638 | 2.210E-04 | Uncharacterized |
| DPOGS211620 | -9.907 | 4.977 | 2.947E-04 | Uncharacterized |
| DPOGS202595 | -9.197 | 4.968 | 3.075E-04 | Serpin-27 |
| DPOGS209028 | -8.462 | 1.370 | 3.075E-04 | Uncharacterized |
| DPOGS207056 | -10.801 | 0.320 | 3.075E-04 | Uncharacterized |
| DPOGS200549 | -5.086 | 1.072 | 3.075E-04 | Aminopeptidase N-like protein |
| DPOGS200623 | -8.542 | 2.970 | 3.075E-04 | Molting fluid carboxypeptidase |

**Table 3.** List of top 15 differentially expressed genes in body tissues between larvae fed with *A. curassavica* and *A. incarnata*. The list includes the top 15 genes significantly up-regulated when <u>fed</u> with *A. curassavica* and the top 15 genes significantly up-regulated when fed with *A. incarnata*.

| Gene ID | log<sub>2</sub>FC | logCPM | FDR | Protein |
| --- | --- | --- | --- | --- |
| ***Top 15 up-regulated genes in A. curassavica*** |  |  |  |  |
| DPOGS202254 | 5.862 | 5.916 | 3.531E-05 | Threonine dehydratase catabolic-like isoform 2 |
| DPOGS207974 | 8.391 | 3.079 | 4.263E-04 | Cuticle protein |
| DPOGS210599 | 5.474 | 4.955 | 5.561E-04 | Cytochrome b5 |
| DPOGS207878 | 7.965 | 6.632 | 6.747E-04 | Antennal binding protein |
| DPOGS209820 | 10.405 | 1.778 | 1.544E-03 | Allantoicase |
| DPOGS204877 | 4.834 | 7.461 | 2.096E-03 | Neuropeptide-like precursor |
| DPOGS209878 | 14.095 | 4.153 | 2.685E-03 | Cuticle protein |
| DPOGS201344 | 4.463 | 4.569 | 2.685E-03 | Uncharacterized |
| DPOGS213427 | 5.256 | 4.346 | 2.893E-03 | Phosphatidyltransferase |
| DPOGS212746 | 4.380 | 10.241 | 3.452E-03 | Uncharacterized |
| DPOGS204901 | 8.429 | 3.785 | 6.396E-03 | Cuticle protein |
| DPOGS202353 | 2.649 | 4.584 | 6.396E-03 | Serine protease inhibitor 32 |
| DPOGS200671 | 9.672 | 2.135 | 6.396E-03 | Cuticle protein |
| DPOGS204876 | 5.325 | 2.603 | 6.911E-03 | Uncharacterized |
| DPOGS204902 | 7.870 | 2.782 | 6.911E-03 | Cuticle protein |
| ***Top 15 up-regulated genes in A. incarnata*** |  |  |  |  |
| DPOGS213127 | -11.298 | 2.225 | 8.082E-06 | Nuclear receptor GRF |
| DPOGS205589 | -10.791 | 5.267 | 1.967E-05 | Hormone receptor 3C |
| DPOGS216089 | -7.901 | 2.515 | 3.531E-05 | Uncharacterized |
| DPOGS209528 | -11.924 | 2.200 | 6.803E-05 | UDP-glycosyltransferase |
| DPOGS207933 | -7.987 | 2.536 | 4.245E-04 | Uncharacterized |
| DPOGS201723 | -8.964 | 3.188 | 4.245E-04 | Peritrophic matrix protein |
| DPOGS209249 | -17.175 | 6.228 | 5.561E-04 | Uncharacterized |
| DPOGS211620 | -12.359 | 5.805 | 1.142E-03 | Uncharacterized |
| DPOGS204937 | -4.721 | 3.391 | 1.358E-03 | Polypeptide N-acetylgalactosaminyltransferase |
| DPOGS212114 | -14.837 | 3.068 | 1.358E-03 | Laccase-like multicopper oxidase 2 |
| DPOGS212041 | -3.204 | 2.933 | 2.685E-03 | Fibroblast growth factor receptor |
| DPOGS207643 | -15.926 | 4.052 | 3.542E-03 | Cytochrome P450 6AB4 |
| DPOGS205455 | -10.749 | 3.336 | 3.542E-03 | Uncharacterized |
| DPOGS213243 | -6.609 | 4.497 | 3.542E-03 | Cytochrome P450 |
| DPOGS201539 | -12.438 | 6.447 | 3.542E-03 | Uncharacterized |

In the body samples, three canonical detoxification genes were up-regulated when fed with *A. incarnata*, including one UDP-glycosyltransferase (DPOGS209528) and two cytochrome P450s (DPOGS207643 and DPOGS213243). In addition, the top 15 up-regulated genes also include a cytochrome b5 (DPOGS210599), which is a redox partner to cytochrome P450 in the P450 system (Després et al., 2007). Five of the top 15 up-regulated genes when fed with *A. curassavica* encode cuticular proteins. Interestingly, cuticle proteins have also been found to be differentially expressed in other insects when feeding on different host plants (*e.g.*, Birnbaum et al., 2017; Celorio-Mancera et al., 2013). Many of the remaining top differentially expressed genes (43.3% in gut and 30.0% in body) have unknown functions.

### 3.4 Examination of specific gene sets

Given existing evidence from other herbivore systems mentioned previously (Smilanich et al., 2009) and our hypothesis that host plants affect immune gene expression, we examined whether any of the known canonical insect immune genes were differentially expressed when feeding on different milkweed species. Among the full set of differentially expressed genes between larvae fed *A. curassavica* and *A. incarnata*, we found that only four immune genes were significantly differentially expressed in gut tissue and only one immune gene was differentially expressed in whole-body tissue (Table 4). For the four differentially expressed immune genes associated with gut samples, two of them are CLIP serine proteases, one is a frep-like receptor, and the other one is a Toll-like receptor. The one differentially expressed gene associated with body samples is a CLIP serine protease that was also differentially expressed in the gut. Interestingly, all four of them were down-regulated in caterpillars fed *A. curassavica*, the more toxic species on which parasite growth was reduced. Overall, we did not find any support that more toxic milkweeds (i.e., *A. curassavica*) enhance the immunity of monarch larvae. Instead, we found weak support that feeding on more toxic milkweeds might cause down-regulation of a subset of immune genes.

**Table 4.**
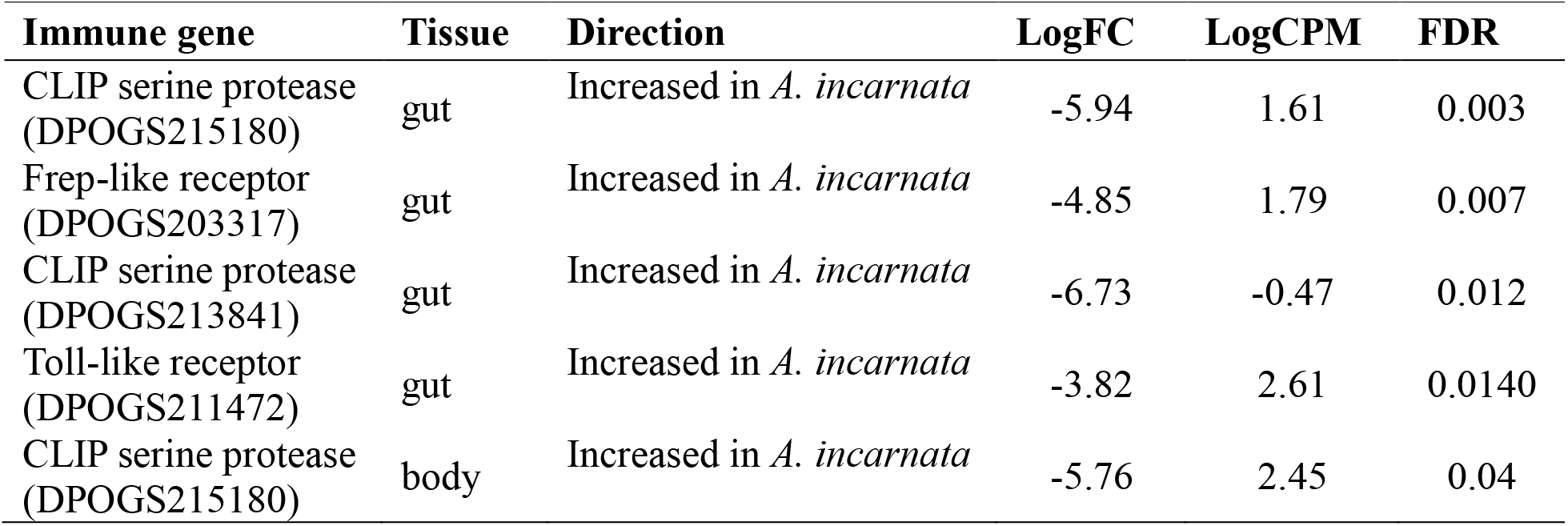
Canonical immune genes that were significantly differentially expressed in gut tissue between larvae fed with *A. curassavica* and *A. incarnata*. No canonical immune genes were significantly differentially expressed between infected and uninfected larvae.

Next, given that monarch larvae were fed with two milkweed species that differ greatly in toxicity, we examined whether any of the known canonical insect detoxification genes were differentially expressed when feeding on the two milkweed species. We focused on gut tissues here because the gut is the place of primary contact with plant materials, where initial digestion and detoxification take place, and because we found stronger differential expression in gut than body tissues. We found that a large proportion of known detoxification genes were expressed (Table 5). Moreover, the proportion of detoxification genes within all significantly differentially expressed genes (2.42%) was significantly higher than the proportion of all annotated genes in the genome that are detoxification genes (1.35%) (χ^2^ = 6.12, *df* = 1, *P* = 0.013), suggesting that they are overrepresented in the genes differentially expressed in monarchs wreared on different milkweeds. The direction of differential expression was not universal, with some genes being up-regulated when on the toxic *A. curassavica* and others when ohen n the less toxic *A. incarnata*. Specifically, 6 CYP450s, 2 UGTs, and 1 GST were up-regulated in monarchs fed *A. curassavica*, while 3 CYP450s, 1 UGTs, 8 ABC transporters, and 1 GST were up-regulated in monarchs fed *A. incarnata* (Table 5 and Supplementary Table S3). Interestingly, all of the ABC transporters were only significantly up-regulated in monarchs fed with *A. incarnata*. Overall, our results demonstrate that several canonical detoxification genes were differentially expressed when larvae fed on the two milkweeds species with different levels of toxicity, suggesting that these genes are involved in metabolizing secondary compounds.

**Table 5.** Canonical detoxification genes that were significantly differentially expressed in gut tissue between larvae fed with *A. curassavica* and *A. incarnata*. The second column, “Annotated”, indicates the number of annotated genes in the genome for the given gene family. The third column, “Expressed”, indicates the number of genes that were expressed in our RNA-seq dataset (defined as counts > 0 in at least two samples). The last two columns show the number of significantly differentially expressed genes.

| Gene family | Annotated | Expressed | Increased in *A. curassavica* | Increased in *A. incarnata* |
| --- | --- | --- | --- | --- |
| Cytochrome P450 (CYP) | 75 | 72 | 6 | 3 |
| UDP glucuronosyltransferases (UGT) | 35 | 34 | 2 | 1 |
| ATP-binding cassette transporters (ABC transporters) | 61 | 60 | 0 | 8 |
| Glutathione S-transferases (GSTs) | 33 | 31 | 1 | 1 |

### 3.5 Gene ontology enrichment analysis

Given that there were almost no differentially expressed genes across infection treatments, we only performed GO enrichment analysis on differentially expressed genes between larvae fed with different plant species. We performed separate analyses for significantly up-regulated genes in larvae fed with *A. curassavica* and significantly up-regulated genes in larvae fed with *A. incarnata* in the two tissue types. Among up-regulated genes in larvae reared on *A. curassavica*, we found a total of 19 GO terms significantly enriched in the gut tissue and one GO term significantly enriched in the body. Among up-regulated genes in *A. incarnata*-reared larvae, we found a total of 112 GO terms significantly enriched in the gut tissue and 6 GO terms significantly enriched in the body (Table 6). Significantly enriched GO terms for each group are shown in Fig. 6 & 7. Overall, we found many more significantly enriched GO terms in gut tissue than in body, and in larvae fed with *A. incarnata.* However, none of those GO terms have seemingly direct functional relevance to detoxification or immunity.

**Figure 6.**
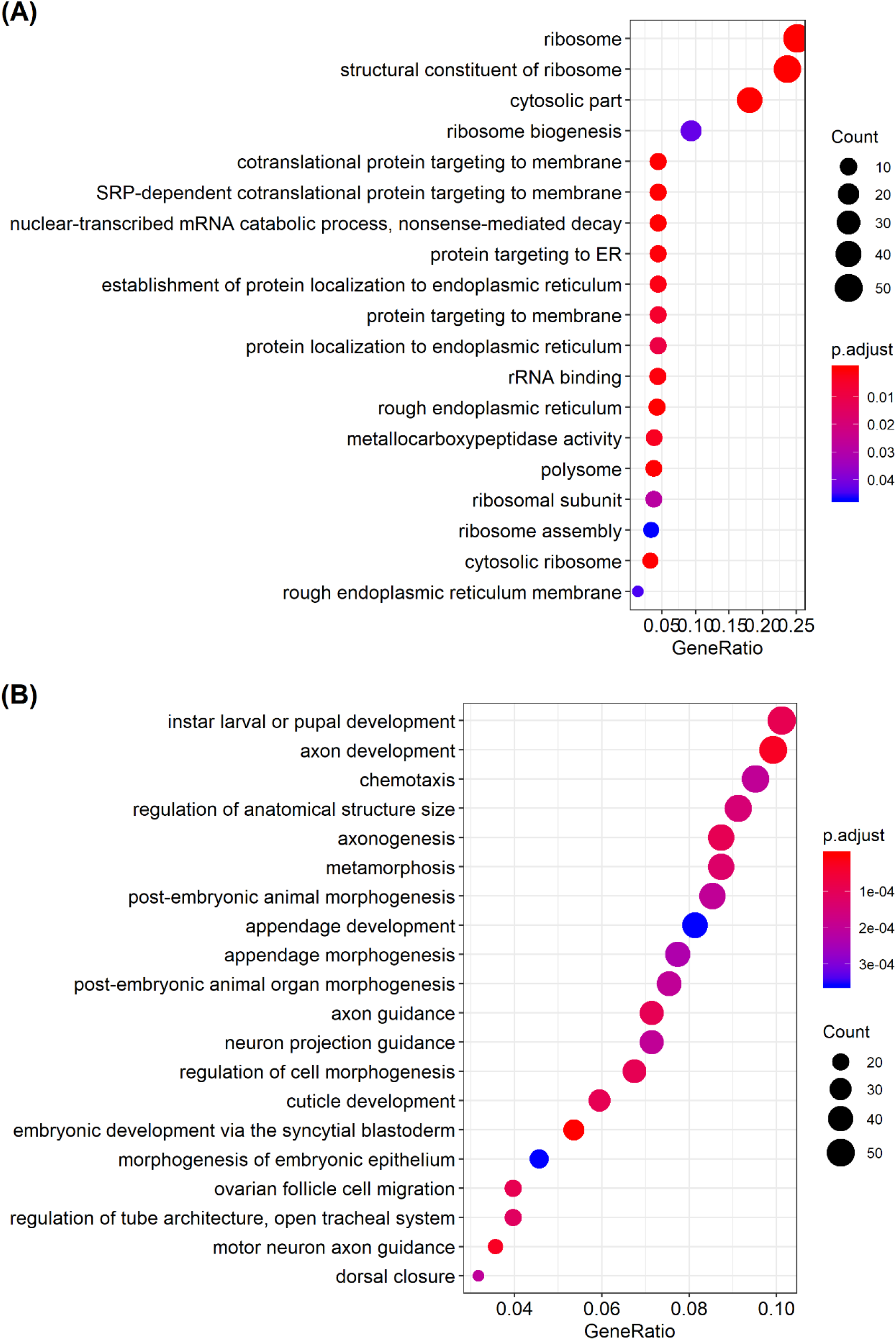
Significantly functionally enriched GO terms in gut tissue between larvae fed with *A. curassavica* and *A. incarnata*. (A) 19 significant terms in up-regulated genes in *A. curassavica*. (B) 116 significant terms in up-regulated genes in *A. incarnata*. Only the top 20 were shown. The x-axis represents the proportion of genes that belong to a given functional category to the total number of differentially expressed genes. All three ontology terms (BP, MF, CC) were included. BP = biological process, MF = molecular function, CC = cellular component. P-values were corrected using the Benjamini-Hochberg method.

**Figure 7.**
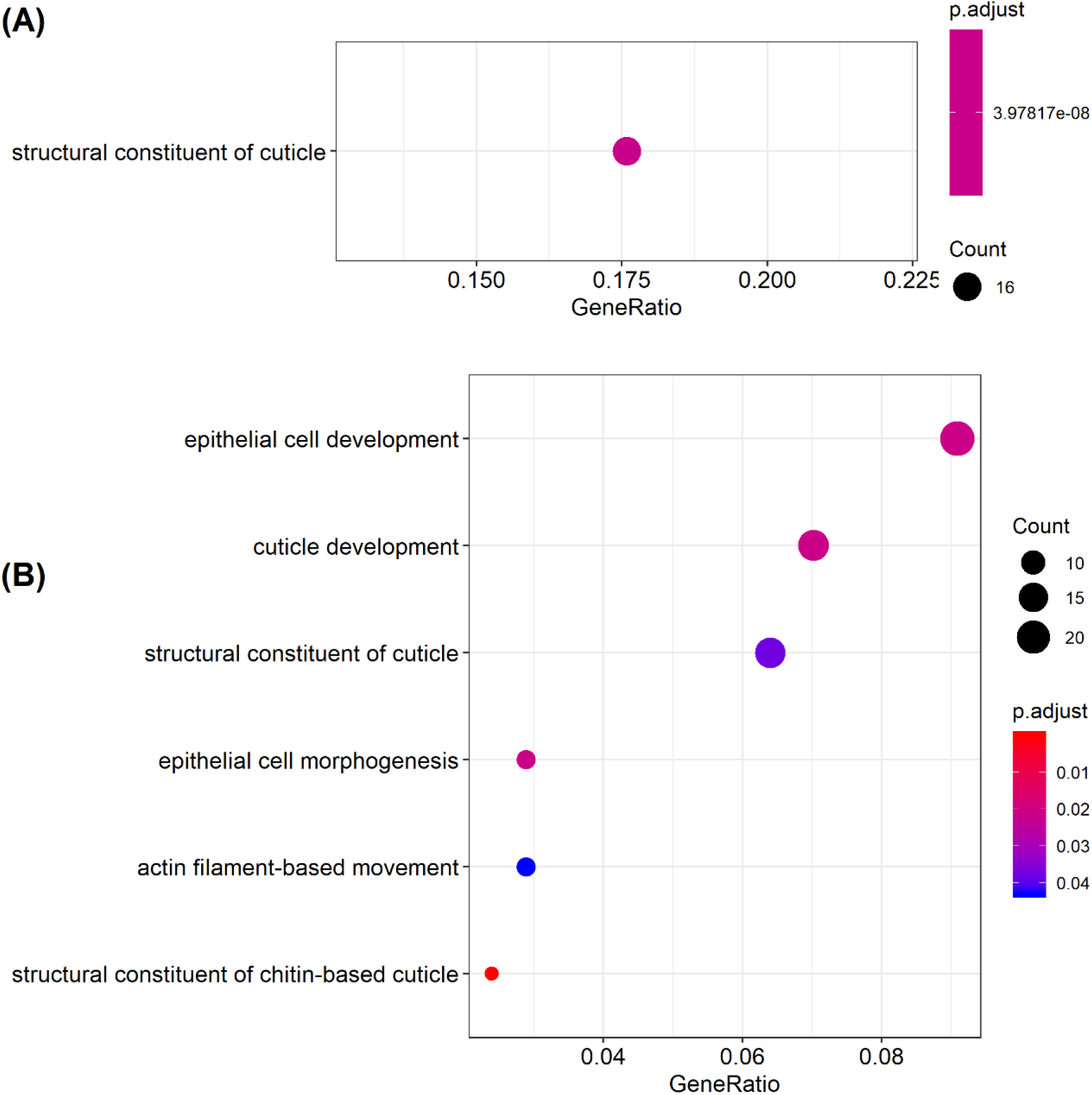
Significantly functionally enriched GO terms in body tissue between larvae fed with *A. curassavica* and *A. incarnata*. (A) One significant term in up-regulated genes in *A. curassavica*. (B) Six significant terms in up-regulated genes in *A. incarnata*. The x-axis represents the proportion of genes that belong to a given functional category to the total number of differentially expressed genes. All three ontology terms (BP, MF, CC) were included. BP = biological process, MF = molecular function, CC = cellular component. P-values were corrected using the Benjamini-Hochberg method.

**Table 6.** Number of significantly functionally enriched GO terms in gut and body tissues between larvae fed with *A. curassavica* and *A. incarnata*. BP = biological process, MF = molecular function, CC = cellular component. Multiple testing was accounted for using the Benjamini-Hochberg method.

| **Tissue type** | **direction** | **BP** | **MF** | **CC** | **Total** |
| --- | --- | --- | --- | --- | --- |
| Gut | Increased in *A. curassavica* | 9 | 3 | 7 | 19 |
| Gut | Increased in *A. incarnata* | 102 | 0 | 10 | 112 |
| Body | Increased in *A. curassavica* | 0 | 1 | 0 | 1 |
| Body | Increased in *A. incarnata* | 4 | 2 | 0 | 6 |

## 4 DISCUSSION

This study examined differences in transcriptional profiles between monarch butterfly larvae feeding on two milkweed species and in response to infection by a specialist protozoan parasite. Our results demonstrate that hundreds of genes were differentially expressed in gut and body when feeding on two different milkweed species. Given that these two milkweed species differ greatly in their concentrations of secondary chemicals (cardenolides) (Fig. 1A) but little in nutrient composition (Tao, Ahmad, et al., 2016), these transcriptional differences are likely related to coping with different levels of toxicity in the diet. Consistent with this hypothesis, we found that several canonical insect detoxification genes were differentially expressed in monarchs reared on the two milkweed species. We discovered that many more genes were differentially expressed in gut than body tissue and that transcriptional profiles of gut samples formed more defined clusters, suggesting that transcriptional responses in relation to milkweed diet are stronger in the gut than in the rest of the body. We also found four canonical immune genes that were differentially expressed between individuals fed on different milkweed species. Interestingly, all four immune genes were down-regulated in monarchs reared on *A. curassavica*, the plant species that reduced parasite infection. In contrast with these transcriptional responses to milkweed diet, we found few transcriptional differences between infected and uninfected monarchs.

### 4.1 Detoxification of plant secondary chemicals

Many plants produce secondary metabolites as defense chemicals against herbivores. In response, herbivorous insects express genes that function in several protective mechanisms, including enzymatic detoxification, excretion, and sequestration (Després et al., 2007). Some previous studies have demonstrated that insects differentially express detoxification genes when feeding on plants with different levels of defense chemicals. For instance, *Drosophila mettleri*, a fruit fly species specialized on cacti with toxic alkaloids, differentially expresses several detoxification genes, including P450s, UGTs, GST, and carboxylesterases, when feeding on different food sources (Hoang, Matzkin, & Bono, 2015). *Tupiocoris notatu*s, a mirid species, down-regulates several GST, UGT, and P450s when feeding on defenseless (JA-silenced) *Nicotiana attenuata* (Crava, Brütting, & Baldwin, 2016). Similarly, our results demonstrate differences in transcriptional profiles of monarch larvae feeding on different milkweed species. Several of those differentially expressed genes belong to canonical detoxification genes, including P450s, UGTs, GSTs, and ABC transporters. Detoxification-related categories, however, were not significantly enriched in our enrichment analyses. While the majority of detoxification genes were expressed, only a relatively small proportion of them were differentially expressed between monarchs reared on the different plant species. Taken together, these results suggest that although a large number of detoxification genes are required for metabolizing a toxic plant diet, only a relatively small proportion of them are related to dealing with variable levels of toxicity. Although our significantly enriched expression categories are not related to detoxification, many of them have also been reported in other studies of herbivorous insects. For instance, categories related to membrane, cuticle, and ribosome are significantly enriched in *Polygonia c-album* when feeding on different plant species (Celorio-Mancera et al., 2013). Enrichment of cuticle-related and developmental-related genes when feeding on different host plants has also been reported in milkweed aphids (Birnbaum et al., 2017) and in several other herbivorous insects (Hoang et al., 2015; Mathers et al., 2017; Matzkin, 2012; Schweizer, Heidel-Fischer, Vogel, & Reymond, 2017; Zhong, Li, Chen, Zhang, & Li, 2017), suggesting that those genes might have pleiotropic effects on detoxification processes, or might be important for structuring of gut tissues. Thickening cuticular components hasbeen suggested to reduce the penetration of insecticides, facilitating insecticide resistance (Foster et al., 2010). Alternatively, as certain insecticides are known to inhibit chitin synthesis (Leighton, Marks, & Leighton, 1981), it is possible that insects regulate the transcription of cuticle-related genes to deal with the interference of plant toxins on chitin metabolism and cuticular protein interactions (Celorio-Mancera et al., 2013).

CYP450 is one of the largest gene families in insects and catalyzes a wide range of reactions (Werck-Reichhart & Feyereisen, 2000). In many insects (e.g., black swallowtail (*Papilio polyxenes*) and parsnip webworm (*Depressaria pastinacella*)), the monooxygenase activity of P450s plays an important role in metabolizing plant toxins such as furanocoumarins (Mao, Rupasinghe, Zangerl, Schuler, & Berenbaum, 2006; Schuler, 1996; Wen, Pan, Berenbaum, & Schuler, 2003). Cardenolides are also substrates for CYP450 monooxygenases (Marty & Krieger, 1984), and it is assumed that milkweed-feeding insects metabolize cardenolides during the detoxification process (Agrawal et al., 2012). Our results indicate that many CYP450 genes are expressed and some of them are differentially expressed when feeding on milkweeds with different levels of cardenolides, suggesting that they play a role in detoxifying cardenolides. Furthermore, our chemical analyses comparing foliage and frass cardenolide composition identified specific cardenolides in frass that are not present in foliage, including several with high polarity. This result, consistent with a recent study (Jones et al., 2019), suggests that some of the cardenolides excreted via frass are likely modified forms, created through detoxification processes. Thus, CYP450 genes may play a role in this modulation, but future studies are needed to directly examine their function.

### 4.2 Specialization on cardenolide-containing plants and sequestration of cardenolides

Despite the fact that milkweed-feeding insects have been one of the most studied systems in chemical ecology and plant-insect interactions, to our knowledge, very few studies have characterized global transcriptional responses of specialist insects when feeding on milkweeds. Recently, Birnbaum et. al. (2017) compared transcriptional profiles using both RNA-seq and qPCR of milkweed aphids (*Aphid nerii*) fed on three different milkweed species, including the plant species used in our study. Similar to our study, they found differential expression of canonical insect detoxification genes, including genes belonging to CYP450s, UGTs, GSTs, and ABC transporters. In addition, their findings and our results both indicate that a greater number of genes are down-regulated rather than up-regulated when milkweed-specialized insects feed on more toxic plant species (Table 1)(Birnbaum et al., 2017). Although both studies on milkweed-feeding insects showed similar results, milkweed aphids do not have the target site mutations on Na^+^/K^+^-ATPase that confer resistance to cardenolides in monarchs (Zhen et al., 2012), suggesting that they rely on other mechanisms to cope with cardenolides. A previous study across three milkweed-feeding butterflies that differ in target site sensitivity indicated that resistance conferred by target site insensitivity has a stronger association with sequestering cardenolides than with digesting cardenolide-rich diets (Petschenka & Agrawal, 2015). Therefore, since the two species differ in target site sensitivity but exhibit similar transcriptional responses to feeding on more toxic plants, the differentially expressed genes may be important in sequestration processes, as both species sequester cardenolides as a defense against predators (Rosenthal & Berenbaum, 1991).

Previous studies have demonstrated that monarch larvae can regulate the level of cardenolide sequestration, as indicated by the fact that cardenolide concentration in larval hemolymph and milkweed leaves do not show a linear relationship (Rosenthal & Berenbaum, 1991). Interestingly, monarchs concentrate cardenolides when feeding on low-cardenolide plants and sequester less when feeding on plants with a very high concentration of cardenolides (Jones et al., 2019; Malcolm, 1991). Notably, our results show that all the differentially expressed ABC transporters were up-regulated in larvae fed *A. incarnata*, a milkweed species with very low cardenolide concentrations. Studies of other insect systems have shown that ABC transporters are involved in sequestration processes. For example, ABC transporters play a key role in salicin sequestration in poplar leaf beetles (*Chrysomela populi*) (Strauss, Peters, Boland, & Burse, 2013). Therefore, the up-regulation of ABC transporters when feeding on low-cardenolide milkweed might be related to an increased rate of cardenolide sequestration.

### 4.3 The effects of plant diet on immunity

Some studies have demonstrated that plant diets with high toxicity can reduce immune responses of herbivorous insects (Smilanich et al., 2009). Detoxification and sequestration of plant toxins can be energetically costly (Bowers, 1992), so a reduction in immune function could be caused by trade-offs with these processes (Moret & Schmid-Hempel, 2000). Plant toxins may have direct negative effects on immune cells (Smilanich et al., 2009). Alternatively, insect hosts may invest less in immunity when anti-parasite resistance is provided by host plants instead. In our study, although we did not find a strong overall effect of plant diet on the expression of canonical immune genes, we observed reduced expression of four immune genes in monarchs feeding on *A. curassavica*, the anti-parasitic plant species. This does not preclude the possibility that other monarch immune defenses not captured by gene expression differences may be influenced by host plant diet. Future studies should couple investigation of immune gene expression with studies of cellular immune responses and should strive to characterize the function of the many genes of unknown function in monarchs, some of which could play a role in anti-parasitic defense.

In the context of herbivore-parasite interactions, medicinal effects conferred by plant diet could be mediated by either direct or indirect effects of plant toxins on parasites. Specifically, medicinal compounds may directly interfere with parasites or may indirectly enhance disease resistance by stimulating immune responses. In the former scenario, investment in immune responses may be reduced because they are compensated for by the medicinal compounds. Indeed, recent studies have demonstrated that the use of medicinal compounds reduces immune investment in a variety of insect species. For example, honey bees (*Apis mellifera*) provided with resins, which have antimicrobial properties, exhibit reduced expression of two immune genes (Simone, Evans, & Spivak, 2009). Similarly, the presence of resins also reduces humoral immune responses in wood ants (*Formica paralugubris*) (Castella, Chapuisat, Moret, & Christe, 2008). Furthermore, long-term association with medicinal compounds might lead to relaxed selection on immune genes. The genome of honey bees (*Apis mellifera*) has a reduced number of canonical insect immune genes, possibly due to the use of medicinal compounds and behavioral defense mechanisms (Evans et al., 2006). Our results show that all four significantly differentially expressed canonical immune genes were down-regulated in monarchs fed with *A. curassavica*, which is in line with the hypothesis that medicinal milkweeds lead to reduced investment in immunity.

Interestingly, one of the immune genes that was down-regulated in larvae feeding on *A. curassavica* is a FREP-like receptor (DPOGS203317). Previous studies of infection of insects by another apicomplexan parasite (*Plasmodium* in *Anopheles gambiae*), which also infects insects through the midgut wall, have shown that several fibrinogen-related proteins (FREPs) play an important role in anti-parasitic defense. For example, overexpression of FREP13 results in increased resistance to *Plasmodium* infection (Dong & Dimopoulos, 2009; Simões et al., 2017). In contrast, inactivation of FREP1 increases resistance, because FREP1 functions as an important host factor that mediates *Plasmodium* ookinete’s invasion of the mosquito midgut epithelium (Dong, Simões, Marois, & Dimopoulos, 2018; Zhang et al., 2015). Our results show down-regulation of a FREP-like gene when larvae feed on a milkweed that confers stronger resistance to parasite infection. However, the exact function of this FREP-like gene remains unknown. In addition, two other immune genes that were down-regulated when feeding on *A. curassavica* are CLIP serine proteases (DPOGS215180 and DPOGS213841). CLIP serine proteases are a large gene family (Christophides et al., 2002), and some of them play an important role in anti-malaria defense (Barillas-Mury, 2007; Volz, Müller, Zdanowicz, Kafatos, & Osta, 2006). Future studies that directly examine the function of these particular immune genes are needed to understand their potential role in defense against *O. elektroscirrha* infections.

### 4.4 Transcriptional responses in relation to parasite infection

Our study confirmed previous findings that monarch larvae fed with *A curassavica* (high-cardenolide) have stronger anti-parasite resistance than those fed with *A. incarnata* (low-cardenolide) (Fig. 1B). Nevertheless, we observed almost no transcriptional response to parasite infection regardless of host plant diet. There are three possible explanations for these results. First, the parasite might be able to suppress or evade the host immune system, which has been demonstrated in several other specialist parasites (Gurung & Kanneganti, 2015; MacGregor, Szöőr, Savill, & Matthews, 2012; Selkirk, Bundy, Smith, Anderson, & Maizels, 2003). Second, the infection may not induce a systemic response; the immune responses may instead have occurred locally and hence may not have been detectable when sequencing the transcriptome of the gut or body. Third, we chose a 24-hr timepoint post infection to try to capture host responses against parasites invading into the body cavity, which is the period in the infection cycle when mosquitoes exhibit up-regulation in midgut-based immune responses to apicomplexan parasites (Blumberg, Trop, Das, & Dimopoulos, 2013; Vlachou, Schlegelmilch, Christophides, & Kafatos, 2005). However, it is possible that the parasite is more active and/or has a stronger interaction with the host immune system at different stages of the infection cycle. Thus, additional life stages should be taken into consideration in future analyses.

## 5 CONCLUSIONS

We compared transcriptional profiles of monarch larvae fed two different milkweed species and examined larval transcriptional responses to infection by a specialist parasite. Our results demonstrate that monarch larvae differentially express hundreds of genes when feeding on *A. curassavica* or *A. incarnata*, two milkweed species that differ strongly in their secondary chemical content. Those differentially expressed genes include genes within multiple families of canonical insect detoxification genes, suggesting that play a role in processing plant diets with different levels of toxicity. Notably, all ABC transporters were up-regulated in monarchs fed with *A. incarnata*, the less toxic plant, which might be related to an increased cardenolide sequestration. Interestingly, the few immune genes that were differentially expressed in monarchs reared on the two plant species were all down-regulated on the anti-parasitic *A. curassavica*, consistent with the hypothesis that medicinal plants could reduce immune investment by providing an alternative form of anti-parasite defense.

## ACKNOWLEDGMENTS

We thank W. Palmer for providing immune gene annotation resources for *Danaus plexippus*, A. Berasategui, E. Diaz-Almeyda, H. Salem, C. Beck, S. Birnbaum, B.-W. Lo, E. Whittington, and V. Talla for helpful discussion on this research, A. Salmi and K. Baffour-Ado for help with the experiments, and H. Streit for performing the chemical analyses. Computational analyses were performed on resources provided by the University of Kansas Information and Telecommunication Technology Center. This work was supported by National Science Foundation (NSF) grant IOS- 1557724 to JCdR, NMG, and MDH, NSF Graduate Research Fellowship Program DGE-1444932 to EVH, and NSF grant DEB-1457758 to JRW.

## DATA ACCESSIBILITY

All sequence data will be archived at the NCBI GeneBank, and other data will be deposited to the Dryad Digital Repository, if the manuscript is accepted for publication. Custom transcriptomic analysis scripts can be found in the following GitHub repository: https://github.com/WaltersLab/Monarch_RNA-Seq

## AUTHOR CONTRIBUTIONS

WHT designed and carried out experiments, performed data analyses, and wrote the initial manuscript. NMG, MDH, and JCdR designed experiments and edited the manuscript. TA, EVH, TYA, and JCdR carried out experiments. JRW provided additional guidance on transcriptomic analyses. MDH supervised chemical analyses. All authors have reviewed and provided comments on the manuscript.

