## Supplemental Information for "Transcriptomics of monarch butterflies (*Danaus plexippus*) reveals strong differential gene expression in response to host plant toxicity, but weak response to parasite infection"

**Table S1.** Sample sizes for each RNA-Seq. treatment group.

| Hostplant species | Infection treatment | Tissue type | Sample size |
|---|---|---|---|
| *A. incarnata* | Infected | Gut | 4 |
| *A. incarnata* | Infected | Body | 4 |
| *A. incarnata* | Uninfected | Gut | 3 |
| *A. incarnata* | Uninfected | Body | 3 |
| *A. curassavica* | Infected | Gut | 3 |
| *A. curassavica* | Infected | Body | 3 |
| *A. curassavica* | Uninfected | Gut | 3 |
| *A. curassavica* | Uninfected | Body | 3 |

**Table S2.** List of immune genes used in this study.

| Gene ID | Gene name | Gene length | Functional class | Chromosome |
|---|---|---|---|---|
| DPOGS200905 | PGRP-like | 3371 | Recognition | Z-chromosome |
| DPOGS209813 | PGRP-like | 1574 | Recognition | Autosome |
| DPOGS206909 | PGRP-like | 1472 | Recognition | Z-chromosome |
| DPOGS206910 | PGRP-like | 7919 | Recognition | Z-chromosome |
| DPOGS207148 | PGRP-like | 15320 | Recognition | Z-chromosome |
| DPOGS209814 | PGRP-like | 2222 | Recognition | Autosome |
| DPOGS206026 | PGRP-like | 3333 | Recognition | Z-chromosome |
| DPOGS212963 | BGRP-like | 7393 | Recognition | Autosome |
| DPOGS215599 | BGRP-like | 2910 | Recognition | Autosome |
| DPOGS212940 | BGRP-like | 4002 | Recognition | Autosome |
| DPOGS212941 | BGRP-like | 3544 | Recognition | Autosome |
| DPOGS212964 | BGRP-like | 3508 | Recognition | Autosome |
| DPOGS212965 | BGRP-like | 3969 | Recognition | Autosome |
| DPOGS203317 | Frep-like | 5328 | Recognition | Z-chromosome |
| DPOGS206045 | Frep-like | 4504 | Recognition | Z-chromosome |
| DPOGS203951 | Frep-like | 10862 | Recognition | Z-chromosome |
| DPOGS210549 | Class B-like SCR | 5148 | Recognition | Autosome |
| DPOGS203180 | Class B-like SCR | 13541 | Recognition | Autosome |
| DPOGS202796 | Class B-like SCR | 9956 | Recognition | Autosome |
| DPOGS214397 | Other SCR | 13196 | Recognition | Autosome |
| DPOGS213636 | Other SCR | 15159 | Recognition | Autosome |
| DPOGS212634 | Other SCR | 14283 | Recognition | Autosome |
| DPOGS202826 | Other SCR | 9352 | Recognition | Autosome |
| DPOGS215836 | TEP-like | 10955 | Recognition | Autosome |
| DPOGS210251 | NIM-like | 13838 | Recognition | Autosome |
| DPOGS210210 | NIM-like | 5318 | Recognition | Autosome |
| DPOGS210211 | NIM-like | 13481 | Recognition | Autosome |
| DPOGS204835 | CLIP-like | 3824 | Modulation | Autosome |
| DPOGS205231 | CLIP-like | 27491 | Modulation | Autosome |
| DPOGS206561 | CLIP-like | 4655 | Modulation | Autosome |
| DPOGS215180 | CLIP-like | 11215 | Modulation | Autosome |
| DPOGS215181 | CLIP-like | 10878 | Modulation | Autosome |
| DPOGS206562 | CLIP-like | 14594 | Modulation | Autosome |
| DPOGS206563 | CLIP-like | 3009 | Modulation | Autosome |
| DPOGS213841 | CLIP-like | 3609 | Modulation | Autosome |
| DPOGS215183 | CLIP-like | 13397 | Modulation | Autosome |
| DPOGS215188 | CLIP-like | 4339 | Modulation | Autosome |
| DPOGS204146 | CLIP-like | 5061 | Modulation | Autosome |

| | | | | |
|---|---|---|---|---|
| DPOGS204147 | CLIP-like | 6895 | Modulation | Autosome |
| DPOGS201678 | CLIP-like | 3492 | Modulation | Autosome |
| DPOGS215220 | CLIP-like | 3818 | Modulation | Autosome |
| DPOGS215098 | CLIP-like | 27161 | Modulation | Autosome |
| DPOGS208169 | CLIP-like | 3911 | Modulation | Autosome |
| DPOGS201966 | CLIP-like | 10337 | Modulation | Autosome |
| DPOGS215182 | CLIP-like | 10004 | Modulation | Autosome |
| DPOGS211355 | CLIP-like | 3601 | Modulation | Autosome |
| DPOGS203664 | CLIP-like | 7166 | Modulation | Autosome |
| DPOGS210568 | CLIP-like | 8283 | Modulation | Autosome |
| DPOGS214570 | CLIP-like | 4377 | Modulation | Autosome |
| DPOGS204148 | CLIP-like | 2905 | Modulation | Autosome |
| DPOGS205210 | CLIP-like | 3342 | Modulation | Autosome |
| DPOGS211237 | CLIP-like | 2943 | Modulation | Autosome |
| DPOGS206224 | CLIP-like | 5720 | Modulation | Autosome |
| DPOGS206217 | CLIP-like | 3300 | Modulation | Autosome |
| DPOGS205206 | CLIP-like | 3310 | Modulation | Autosome |
| DPOGS209809 | SPZ-like | 1777 | Signaling - Toll | Autosome |
| DPOGS209810 | SPZ-like | 5018 | Signaling - Toll | Autosome |
| DPOGS203200 | Toll_like-receptors | 3821 | Signaling - Toll | Autosome |
| DPOGS205279 | Toll_like-receptors | 4664 | Signaling - Toll | Autosome |
| DPOGS202626 | Toll_like-receptors | 2709 | Signaling - Toll | Autosome |
| DPOGS205281 | Toll_like-receptors | 3894 | Signaling - Toll | Autosome |
| DPOGS205295 | Toll_like-receptors | 3887 | Signaling - Toll | Autosome |
| DPOGS205123 | Toll_like-receptors | 5897 | Signaling - Toll | Autosome |
| DPOGS211472 | Toll_like-receptors | 4533 | Signaling - Toll | Autosome |
| DPOGS200002 | Toll_like-receptors | 5428 | Signaling - Toll | Autosome |
| DPOGS205283 | Toll_like-receptors | 1949 | Signaling - Toll | Autosome |
| DPOGS215274 | Toll_like-receptors | 5667 | Signaling - Toll | Autosome |
| DPOGS203198 | Toll_like-receptors | 3410 | Signaling - Toll | Autosome |
| DPOGS205293 | Toll_like-receptors | 869 | Signaling - Toll | Autosome |
| DPOGS205296 | Toll_like-receptors | 2006 | Signaling - Toll | Autosome |
| DPOGS207788 | Tollip | 3828 | Signaling - Toll | Autosome |
| DPOGS205936 | MyD88 | 2965 | Signaling - Toll | Autosome |
| DPOGS208945 | Tube | 2121 | Signaling - Toll | Autosome |
| DPOGS214647 | Pellino | 3601 | Signaling - Toll | Autosome |
| DPOGS210260 | Pelle | 11577 | Signaling - Toll | Autosome |
| DPOGS202662 | TRAF2 | 6521 | Signaling - Toll | Autosome |
| DPOGS209243 | ECSIT | 1486 | Signaling - Toll | Autosome |
| DPOGS209453 | Cactus | 2181 | Signaling - Toll | Autosome |

| | | | | |
|---|---|---|---|---|
| DPOGS215778 | IMD | 1319 | Signaling - IMD | Autosome |
| DPOGS200403 | TAK1 | 12371 | Signaling - IMD | Autosome |
| DPOGS202907 | IKKgamma | 14063 | Signaling - IMD | Autosome |
| DPOGS202564 | IKKbeta | 1586 | Signaling - IMD | Autosome |
| DPOGS207960 | FADD | 916 | Signaling - IMD | Autosome |
| DPOGS212093 | Dredd | 1508 | Signaling - IMD | Autosome |
| DPOGS200977 | Tab2 | 3087 | Signaling - IMD | Autosome |
| DPOGS203759 | IAP2 | 5539 | Signaling - IMD | Autosome |
| DPOGS201405 | Ubc13 | 455 | Signaling - IMD | Autosome |
| DPOGS208954 | Hem | 3377 | Signaling - JNK | Autosome |
| DPOGS213169 | JNK | 3769 | Signaling - JNK | Autosome |
| DPOGS214573 | Fos | 2706 | Signaling - JNK | Autosome |
| DPOGS202887 | Jun | 242 | Signaling - JNK | Autosome |
| DPOGS214325 | PIAS | 12079 | Signaling - JAK-STAT | Autosome |
| DPOGS214451 | SOCS | 2617 | Signaling - JAK-STAT | Autosome |
| DPOGS200349 | DOMELESS | 2960 | Signaling - JAK-STAT | Autosome |
| DPOGS210157 | Hopscotch | 8831 | Signaling - JAK-STAT | Autosome |
| DPOGS212956 | Stat | 16032 | Signaling - JAK-STAT | Autosome |
| DPOGS213997 | Attacin-Like | 877 | Effector | Autosome |
| DPOGS205720 | Attacin-Like | 1439 | Effector | Autosome |
| DPOGS215451 | Attacin-Like | 818 | Effector | Autosome |
| DPOGS210270 | Cecropin-like | 374 | Effector | Autosome |
| DPOGS210268 | Cecropin-like | 422 | Effector | Autosome |
| DPOGS210269 | Cecropin-like | 529 | Effector | Autosome |
| DPOGS200256 | Cecropin-like | 428 | Effector | Autosome |
| DPOGS210271 | Cecropin-like | 352 | Effector | Autosome |
| DPOGS210265 | Cecropin-like | 848 | Effector | Autosome |
| DPOGS210304 | Gloverin-like | 644 | Effector | Autosome |
| DPOGS210303 | Gloverin-like | 822 | Effector | Autosome |
| DPOGS202093 | NOS-like | 15626 | Effector | Autosome |
| DPOGS202094 | NOS-like | 19723 | Effector | Autosome |
| DPOGS201818 | PPO-like | 4660 | Effector | Z-chromosome |
| DPOGS201819 | PPO-like | 5674 | Effector | Z-chromosome |
| DPOGS206820 | PPO-like | 9352 | Effector | Z-chromosome |
| DPOGS200017 | PPO-like | 5087 | Effector | Autosome |

(A)

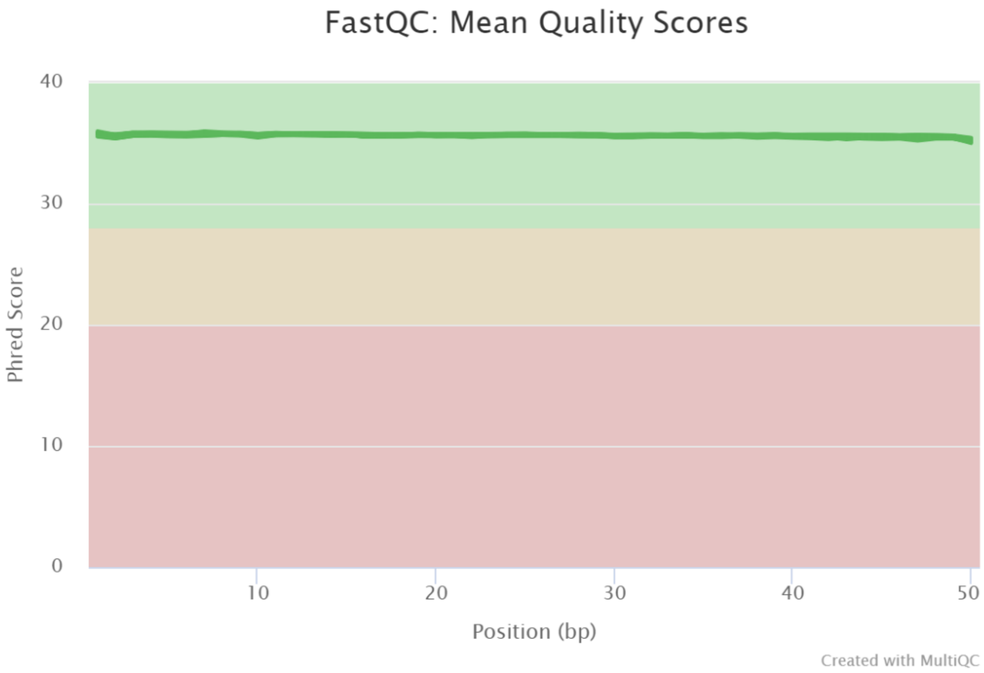


(B)

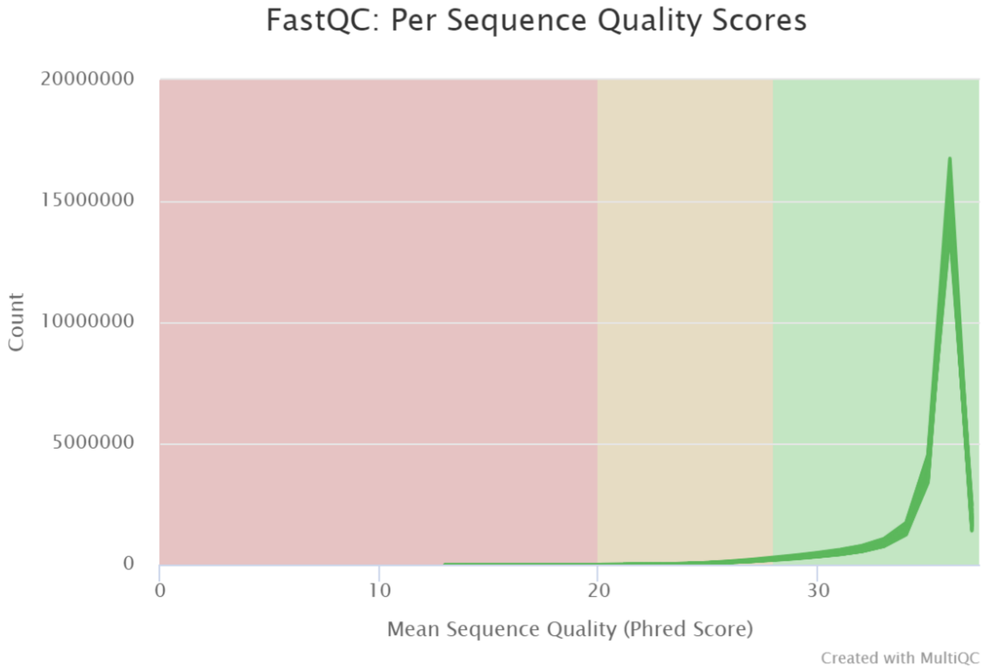


**Figure S1.** Quality of RNA-Seq reads across all samples. (A) The mean quality score across each base position in the read. (B) The number of reads with each mean quality score. Each green curve represents one RNA-Seq sample (N = 26). The curves are highly overlapped across samples because of highly consistent quality.
